# Gene model for the ortholog of *raptor* in *Drosophila grimshawi*

**DOI:** 10.64898/2026.07.07.737051

**Authors:** Bethany C. Lieser, Bailey Lose, Cole A. Kiser, Sydney Butterfield, Luke Laschober, Leon F. Laskowski, Jonas Nielsen, John Pulford, Jeffrey S. Thompson, Chinmay P. Rele, Jacqueline K. Wittke-Thompson

## Abstract

Gene model for the ortholog of *raptor* in the D. grimshawi May 2011 (Agencourt dgri_caf1/DgriCAF1) Genome Assembly (GenBank Accession: GCA_000005155.1) of *Drosophila grimshawi*. This ortholog was characterized as part of a developing dataset to study the evolution of the Insulin/insulin-like growth factor signaling pathway (IIS) across the genus *Drosophila* using the Genomics Education Partnership gene annotation protocol for Course-based Undergraduate Research Experiences.

## Introduction

*This article reports a predicted gene model generated by undergraduate work using a structured gene model annotation protocol defined by the Genomics Education Partnership (GEP; thegep.org) for Course-based Undergraduate Research Experience (CURE). The following information in quotes may be repeated in other articles submitted by participants using the same GEP CURE protocol for annotating Drosophila species orthologs of Drosophila melanogaster genes in the insulin signaling pathway*.

“In this GEP CURE protocol students use web-based tools to manually annotate genes in non-model *Drosophila* species based on orthology to genes in the well-annotated model organism fruit fly *Drosophila melanogaster*. The GEP uses web-based tools to allow undergraduates to participate in course-based research by generating manual annotations of genes in non-model species (Rele et al., 2023). Computational-based gene predictions in any organism are often improved by careful manual annotation and curation, allowing for more accurate analyses of gene and genome evolution (Mudge and Harrow 2016; Tello-Ruiz et al., 2019). These models of orthologous genes across species, such as the one presented here, then provide a reliable basis for further evolutionary genomic analyses when made available to the scientific community.” (Myers et al., 2024).

“The particular gene ortholog described here was characterized as part of a developing dataset to study the evolution of the Insulin/insulin-like growth factor signaling pathway (IIS) across the genus *Drosophila*. The Insulin/insulin-like growth factor signaling pathway (IIS) is a highly conserved signaling pathway in animals and is central to mediating organismal responses to nutrients (Hietakangas and Cohen 2009; Grewal 2009).” (Myers et al., 2024).

“raptor positively regulates (Target of Rapamycin) TOR-mediated cell apoptosis and growth control by differentially regulating S6K-dependent signaling pathways, and is a crucial regulator of cell growth and metabolism in *Drosophila* (Lee and Chung 2007; Hatfield et al., 2015). *raptor* is orthologous to the human *RPTOR* gene (*regulatory associated protein of raptor complex 1*) and is a well conserved component of the TORC1 complex (Wang et al., 2012).” (Backlund et al., 2025).

*“D. grimshawi* is a member of the Picture Wing clade (*sensu* Kaneshiro et al., 1995) of the Hawaiian *Drosophila*. Molecular analyses place the monophyletic Hawaiian *Drosophila* as sister to Scaptomyza clade, which both are nested within the *Drosophila* subgenus of the genus *Drosophila* (Kambysellis et al., 1995; Baker and DeSalle 1997). The Picture Wings are so called due to their dramatically pigmented wings. *D. grimshawi* was first described by Oldenberg (1914), and is found in high elevation cool tropical rainforest on the Maui Complex islands where they breed on rotting vegetation (Carson et al., 1970; Carson 1983).” (Lawson et al., 2024).

We propose a gene model for the *D. grimshawi* ortholog of the *D. melanogaster* raptor (*raptor*) gene. The genomic region of the ortholog corresponds to the uncharacterized protein XP_043072238.1 (Locus ID LOC6566000) in the D. grimshawi May 2011 (Agencourt dgri_caf1/DgriCAF1) Genome Assembly of *D. grimshawi* (GCA_000005155.1). This model is based on RNA-Seq data from *D. grimshawi* (SRP073087) and *raptor* in *D. melanogaster* using FlyBase release FB2024_02 (GCA_000001215.4; Gramates et al., 2022; Jenkins et al., 2022; Larkin et al., 2021).

### Synteny

The reference gene, *raptor*, occurs on chromosome X in *D. melanogaster* and is nested by *Multiple inositol polyphosphate phosphatase 2* (*Mipp2*), flanked upstream by *Ca2+-channel protein* α*1 subunit T* (*Ca-*α*1T*) and *Neprilysin 1* (*Nep1*), and flanked downstream by *CG4660* and *CG4666*. The *tblastn* search of *D. melanogaster* raptor-PC (query) against the *D. grimshawi* (GenBank Accession: GCA_000005155.1 Genome Assembly (database) placed the putative ortholog of *raptor* within scaffold scaffold_14853 (CH916371.1) within locus LOC6566000 (XP_043072238.1)— with an E-value of 0.0 and a percent identity of 61.22%. Furthermore, the putative ortholog is nested by LOC6566000 (XP_001992616.1) and flanked upstream by LOC6566003 (XP_001992619.1), LOC6566002 (XP_032594964.1), and LOC6566001 (XP_001992617.2), which correspond to *Mipp2, Major Facilitator Superfamily Transporter 10* (*MFS10*), *Nep1*, and *CG33107* in *D. melanogaster* (E-value: 0.0, 0.0, 0.0, and 8e-31; identity: 72.26%, 75.95%, 81.40%, and 29.75%, respectively, as determined by *blastp*; Figure 1A, Altschul et al., 1990). The *raptor* and *Mipp2* orthologs share the same LOCID (LOC6566000) but have different protein IDs because they exist within the same locus in the scaffold yet have different predicted coding sequences. The putative ortholog of *raptor* is flanked downstream by LOC6565999 (XP_001992615.1) and LOC6565998 (XP_001992614.1), which correspond to *CG4660* and *CG4666* in *D. melanogaster* (E-value: 1e-136 and 1e-124; identity: 67.14% and 85.49%, respectively, as determined by *blastp*). The putative ortholog assignment for *raptor* in *D. grimshawi* is supported by the following evidence: Although the upstream neighborhood is partially conserved, the genes downstream of the *raptor* ortholog are orthologous to the genes at the same locus in *D. melanogaster* and local synteny is completely conserved, supported by E-values and percent identities, so we conclude that LOC6566000 is the correct ortholog of *raptor* in *D. grimshawi* (Figure 1A).

**Figure 1.**
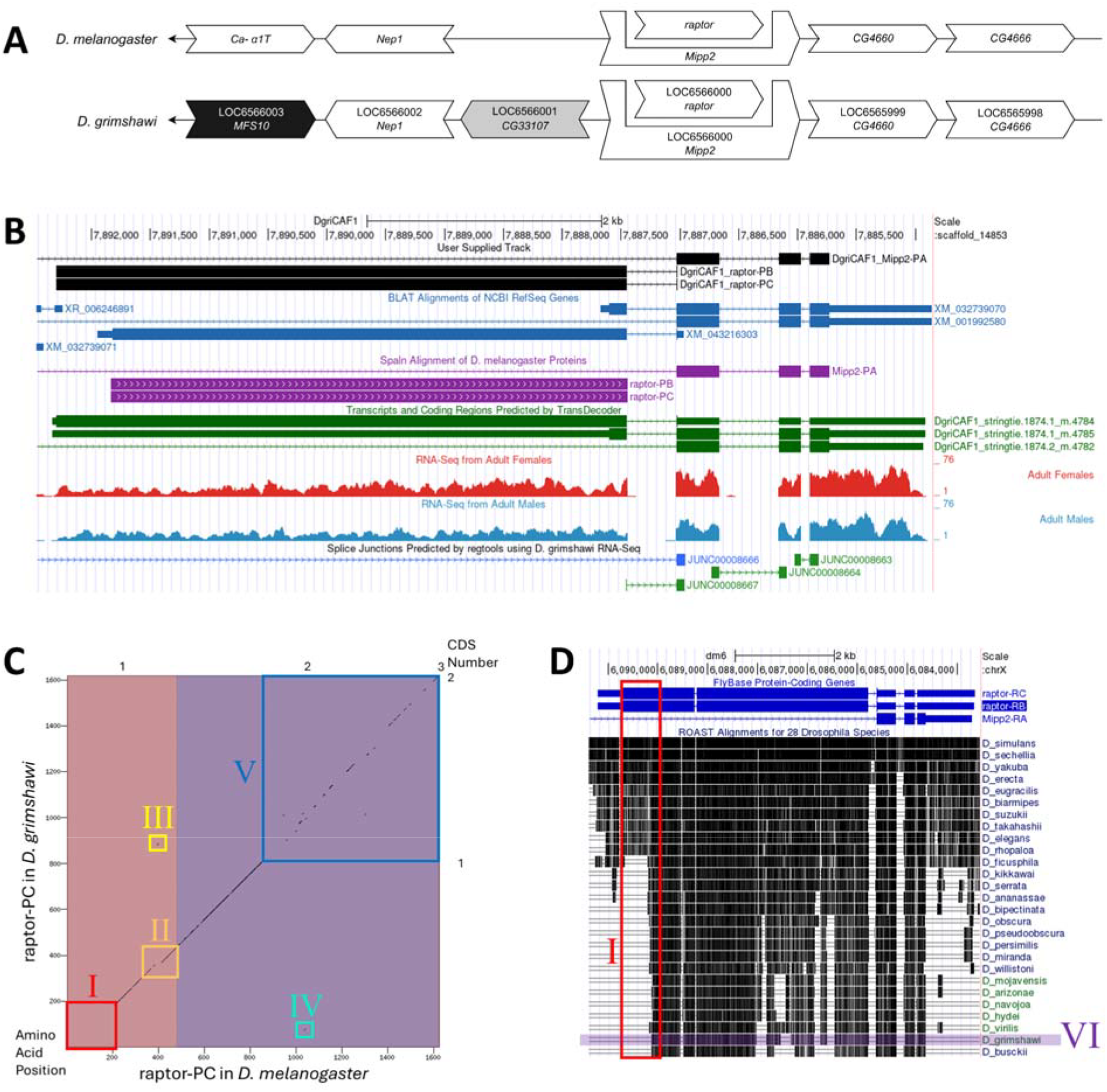
*raptor* gene model comparison between *Drosophila grimshawi* and *Drosophila melanogaster* orthologs. **(A) Synteny comparison of the genomic neighborhoods for *raptor* in *Drosophila melanogaster* and *D. grimshawi***. Thin underlying arrows indicate the DNA strand within which the target gene–*raptor*–is located in *D. melanogaster* (top) and *D. grimshawi* (bottom). The thin arrows pointing to the left indicate that *raptor* is on the negative (-) strand in *D. melanogaster* and *D. grimshawi*. The wide gene arrows pointing in the same direction as *raptor* are on the same strand relative to the thin underlying arrows, while wide gene arrows pointing in the opposite direction of *raptor* are on the opposite strand relative to the thin underlying arrows. White gene arrows in *D. grimshawi* indicate orthology to the corresponding gene in *D. melanogaster*, black gene arrows indicate non-orthology, and grey gene arrows indicate a gene insertion. Gene symbols given in the *D. grimshawi* gene arrows indicate the orthologous gene in *D. melanogaster*, while the locus identifiers are specific to *D. grimshawi*. **(B) Gene Model in GEP UCSC Track Data Hub (Raney et al., 2014)**. The coding-regions of *raptor* in *D. grimshawi* are displayed in the User Supplied Track (black); coding CDSs are depicted by thick rectangles and introns by thin lines with arrows indicating the direction of transcription. Subsequent evidence tracks include BLAT Alignments of NCBI RefSeq Genes (dark blue, alignment of Ref-Seq genes for *D. grimshawi*), Spaln of D. melanogaster Proteins (purple, alignment of Ref-Seq proteins from *D. melanogaster*), Transcripts and Coding Regions Predicted by TransDecoder (dark green), RNA-Seq from Adult Females and Adult Males (red and light blue, respectively; alignment of Illumina RNA-Seq reads from *D. grimshawi*), and Splice Junctions Predicted by regtools using *D. grimshawi* RNA-Seq (SRP073087). The splice junction supporting *raptor* (JUNC00008667), shown in green, has a read-depth of 64. **(C) Dot Plot of raptor-PC in *D. melanogaster* (*x*-axis) vs. the orthologous peptide in *D. grimshawi* (*y*-axis)**. Amino acid number is indicated along the left and bottom; CDS number is indicated along the top and right, and CDSs are also highlighted with alternating colors. Line breaks in the dot plot indicate mismatching amino acids at the specified location between species. The red box denoted I highlights an area of sequence dissimilarity between *D. grimshawi* and *D. melanogaster*, the orange box denoted II outlines two indels, and the yellow and green boxes denoted III and IV highlight two different repeats, ‘SGG’ and ‘GATA’, respectively. The blue box denoted V outlines a region with low sequence conservation. **(D) Conservation of *raptor* across 28 *Drosophila* species shown in the *D. melanogaster* UCSC Genome Browser**. The red box denoted I highlights the beginning of the first CDS of *raptor* in *D. melanogaster* and the sequence conservation of this region in 28 *Drosophila* species, listed along the right, relative to *D. melanogaster*. The black boxes and lines indicate conservation. The sequence of *D. grimshawi* in this region is highlighted by the transparent purple box denoted VI.

### Protein Model

*raptor* in *D. grimshawi* has two CDSs within the genome sequence. The protein sequence (raptor-PB and raptor-PC) is translated from two mRNA isoforms that differ in their UTRs (*raptor-RB* and *raptor-RC*; Figure 1B). Relative to the ortholog in *D. melanogaster*, the protein isoform count is conserved but the CDS number is not conserved. In *D. melanogaster*, each mRNA isoform of *raptor* contains three CDSs. The sequence of raptor-PC in *D. grimshawi* has 65.67% identity (E-value: 0.0) with the protein-coding isoform raptor-PC in *D. melanogaster*, as determined by *blastp* (Figure 1C). Unusual characteristics of this model include the lack of sequence similarity present in CDS one. Coordinates of this curated gene model (*raptor-RB* and *raptor-RC*) are stored by NCBI at GenBank/BankIt (accession **<u>BK065261</u>** and **<u>BK065262</u>**, respectively). This gene model can also be seen within the target genome at this <u>TrackHub</u>.

### Special characteristics of the protein model

#### Lack of sequence similarity in CDS one

Red box I in the Dot Plot outlines a large region of CDS one which lacks sequence similarity to *D. melanogaster*. This region is also outlined by the red box denoted I in Figure 1D, where the conservation of *D. grimshawi* at this location is highlighted by transparent purple box V. The intersection of red box I and transparent purple box VI displays a lack of conservation in *D. grimshawi* in this region. Additionally, a lack of sequence similarity in this region can be seen in seventeen other species (Figure 1D).

## Methods

“Detailed methods including algorithms, database versions, and citations for the complete annotation process can be found in Rele et al. (2023). Briefly, students use the GEP instance of the UCSC Genome Browser v.435 (https://gander.wustl.edu; Kent et al., 2002; Navarro Gonzalez et al., 2021) to examine the genomic neighborhood of their reference IIS gene in the *D. melanogaster* genome assembly (Aug. 2014; BDGP Release 6 + ISO1 MT/dm6). Students then retrieve the protein sequence for the *D. melanogaster* reference gene for a given isoform and run it using *tblastn* against their target *Drosophila* species genome assembly on the NCBI BLAST server (https://blast.ncbi.nlm.nih.gov/Blast.cgi; Altschul et al., 1990) to identify potential orthologs. To validate the potential ortholog, students compare the local genomic neighborhood of their potential ortholog with the genomic neighborhood of their reference gene in *D. melanogaster*. This local synteny analysis includes at minimum the two upstream and downstream genes relative to their putative ortholog. They also explore other sets of genomic evidence using multiple alignment tracks in the Genome Browser, including BLAT alignments of RefSeq Genes, Spaln alignment of *D. melanogaster* proteins, multiple gene prediction tracks (e.g., GeMoMa, Geneid, Augustus), and modENCODE RNA-Seq from the target species. Detailed explanation of how these lines of genomic evidenced are leveraged by students in gene model development are described in Rele et al. (2023). Genomic structure information (e.g., CDSs, intron-exon number and boundaries, number of isoforms) for the *D. melanogaster* reference gene is retrieved through the Gene Record Finder (https://gander.wustl.edu/~wilson/dmelgenerecord/index.html; Rele et al., 2023). Approximate splice sites within the target gene are determined using *tblastn* using the CDSs from the *D. melanogaste*r reference gene. Coordinates of CDSs are then refined by examining aligned modENCODE RNA-Seq data, and by applying paradigms of molecular biology such as identifying canonical splice site sequences and ensuring the maintenance of an open reading frame across hypothesized splice sites. Students then confirm the biological validity of their target gene model using the Gene Model Checker (https://gander.wustl.edu/~wilson/dmelgenerecord/index.html; Rele et al., 2023), which compares the structure and translated sequence from their hypothesized target gene model against the *D. melanogaster* reference gene model. At least two independent models for a gene are generated by students under mentorship of their faculty course instructors. Those models are then reconciled by a third independent researcher mentored by the project leaders to produce the final model. Note: comparison of 5’ and 3’ UTR sequence information is not included in this GEP CURE protocol.” (Gruys et al., 2025)

## Supporting information

Gene model data files

## Supplemental Files

1. Zip file containing a FASTA, PEP, GFF files for the gene model
2. Figure 1 in high resolution

## Metadata

Bioinformatics, Genomics, *Drosophila*, Genotype Data, New Finding

## Acknowledgements

We would like to thank Wilson Leung for developing and maintaining the technological infrastructure that was used to create this gene model and Laura K. Reed for overseeing the project. Thank you to <u>FlyBase</u> for providing the definitive database for *Drosophila melanogaster* gene models. Further, we would like to thank the editors and developers at the journal *microPublication: Biology* for assistance in developing the template for these single gene ortholog publications.

## Funding

This material is based upon work supported by the National Science Foundation (1915544) and the National Institute of General Medical Sciences of the National Institutes of Health (R25GM130517) to the Genomics Education Partnership (GEP; https://thegep.org/; PI-LKR). Any opinions, findings, and conclusions or recommendations expressed in this material are solely those of the author(s) and do not necessarily reflect the official views of the National Science Foundation nor the National Institutes of Health.

## Notes

https://gander.wustl.edu/cgi-bin/hgTracks?db=DgriCAF1&lastVirtModeType=default&lastVirtModeExtraState=&virtModeType=default&virtMode=0&nonVirtPosition=&position=scaffold_14853%3A7886521-7892799&hgct_customText=track%20type=bigGenePred%20visibility=pack%20bigDataUrl=http://genemodels01.ua.edu/trackhub_hosting/models/DgriCAF1/DgriCAF1.bb

### Competing Interest Statement

The authors have declared no competing interest.

